# Spontaneous mutant in threespine stickleback connects endosome trafficking disorders and inflammatory bowel diseases via changes in the gut microbiome

**DOI:** 10.1101/2025.09.20.677535

**Authors:** Emily A. Beck, Caitlin A. Smith, Mark C. Currey, Clayton M. Small, Brendan J. M. Bohannan, William A. Cresko

**Author notes:** Authors contributed equally (Emily A. Beck, Caitlin A. Smith). **Correspondence**: E.A.B. & W.A.C.

## Abstract

Endosomes are organelles that sort and traffic materials to different cellular compartments to enable performance of other organelles. One group of organelles supported by endosomes are lysosome related organelles (LROs) – which perform specialized functions using endocytosed or recycled materials from endosomes. LROs include melanosomes, lytic granules, platelet dense granules, lamellar bodies, and more. Individuals with endosome trafficking disorders commonly present with oculocutaneous albinism due to insufficient trafficking to – and maturation of – melanosomes. While the loss of pigment is an easily identifiable phenotype, mutations in endosome trafficking can also cause multi-organ system symptoms due to more broad disruption of LROs. Different endosome trafficking mutations impact LRO maturation at different degrees of severity and can impact a subset or all LROs leading to a wide range of symptomologies. Endosome trafficking disorders can present with oculocutaneous albinism (caused by mutated melanosomes), blood diathesis (caused by mutated platelet dense granules), pulmonary fibrosis (caused by mutated lamellar bodies), or chronic inflammation and inflammatory bowel disease (IBD) (caused by mutated lytic granules), as well as other symptoms. The shared inflammatory patterns of Hermansky-Pudlak Syndrome (HPS) and Inflammatory Bowel Disease (IBD) are well studied. However, impacts of this shared gastrointestinal inflammation on the gut microbiome remain unexplored. Using a spontaneously identified mutant family of threespine stickleback fish, we characterize changes in LROs and symptoms of oculocutaneous albinism. We further demonstrate that the shared gastrointestinal phenotypes of HPS and IBD extend to the gut microbiome showing similar shifts in microbial communities consistent with IBD. We argue endosome trafficking models should be explored as models to study the gut microbiome in addition to inflammatory phenotypes in IBD, and importantly, be used to test microbial/dietary interventions for both IBD and endosome trafficking disorders.

## INTRODUCTION

Endosomes act as hubs of cellular trafficking, sorting endocytosed materials from the plasma membrane for degradation by the endo-lysosomal pathway or recycling it through various recycling pathways (Jovic et al. 2009; Naslavsky and Caplan 2018). Many specialized organelles rely on endosome trafficking for maturation and function including lysosome-related organelles (LROs) like melanosomes, platelet dense granules, lytic granules, lung lamellar bodies, and more (Bowman et al. 2019). Each of these organelle types rely on materials from recycling endosomes to perform their functions (Marks et al. 2013). Without proper endosome trafficking LROs can be disrupted which can lead to a wide range of impacts on cellular physiology (Marks et al. 2013). In humans, the failure LROs to mature properly can therefore result in platelet storage and lysosomal storage diseases, as well as a group of syndromes called Hermansky-Pudlak Syndromes (HPS) (Dell’Angelica et al. 1999; Bowman et al. 2019), characterized by oculocutaneous albinism, vision impairment, blood diathesis, pulmonary fibrosis, granulomatous colitis, and immunodeficiency (Wei 2006; Hurford and Sebastiano 2008; Bowman et al. 2019; Introne et al. 2023). The loss of pigmentation is caused by loss of melanosomes while blood clotting deficiencies and blood diathesis are caused by a loss of platelet dense granules and Weibel-Palade bodies in endothelial cells. The cause of pulmonary fibrosis is not fully understood, but is associated in part by a loss of lung lamellar bodies impacting surfactant secretion combined with impacts on the immune system caused by deficiencies in many types of LROS loss including lytic granules and IRF7 signaling lysosomes (Huizing et al. 2008; Bowman et al. 2019).

The severity and combination of symptoms associated with endosome trafficking disorders depend on which stage(s) of endosome trafficking are perturbed (Wei 2006; Introne et al. 2023). There are several genetic mutations in different genes that have been previously identified in individuals with endosome trafficking disorders each with their own set of symptoms. In less severe types of HPS, such as HPS Type 5 caused by a mutation in *hps5*, patients present with albinism and mild vision loss but do not exhibit severe symptoms impacting other organ-systems and show no signs of pulmonary fibrosis. In more severe types of HPS, such as HPS Type 4 caused by a mutation in *hps4*, symptoms are much more severe and include a life expectancy at 40-50 years with the majority of patients dying from complications related to the disease (Introne et al. 2023).

In humans with severe HPS, pulmonary fibrosis is the leading cause of mortality. The combined loss of surfactant secretion in the lungs, chronic inflammation, and disruptions to the immune system lead to lung damage – though the full genetic and cellular basis of pulmonary fibrosis is not understood (Huizing et al. 2008; Yokoyama and Gochuico 2021). Chronic inflammation in HPS patients is not exclusive to the lungs and can also impact the gastrointestinal system, leading to inflammatory bowel disease (IBD) (O’Brien et al. 2021; Introne et al. 2023). Chronic inflammation, especially in the bowel, while not the leading cause of death in humans with endosome trafficking disorders can be a life-altering symptom requiring intervention (O’Brien et al. 2021). The loss of platelet dense granules and Weibel-Palade bodies in some HPS patients lead to clotting disorders which contraindicates surgical interventions like bowel resections to treat colitis or lung transplants to treat pulmonary fibrosis. However, interventions including platelet transfusions have worked in many to allow for successful resection or transplant (Van Dorp et al. 1990; Lederer et al. 2005; El-Chemaly et al. 2018). Still, development of non-surgical interventions is needed.

Shared HPS symptoms of colitis are consistent with Inflammatory Bowel Diseases (IBD) (Schinella et al. 1980; Erzin et al. 2006; Grucela et al. 2006; Hazzan et al. 2006; Leusse et al. 2006; Wei 2006; Kouklakis et al. 2007). However, current connections between gut microbiome shifts in patients with IBD and endosome trafficking disorders are unexplored. Host-microbe interactions play an important role in regulating host health and preventing colonization of pathogens (Chung et al. 2012; Pickard et al. 2017). We also know that dietary interventions – including transition to a low inflammatory diet – as well as microbial interventions – including fecal transplants – have been successful in partially mitigating negative effects of IBD (Oka and Sartor 2020). More recently, benefits of anti-inflammatory diets have been shown in not just gastrointestinal symptoms, but seemingly unrelated organ-systems like the lungs (Wood et al. 2015). We argue dietary or microbial interventions may be fruitful avenues to explore for the development of non-surgical therapeutics to HPS patients. First, microbial connections between IBD and endosome trafficking disorders must be established.

We serendipitously discovered a group of mutants in an F2 family of threespine stickleback fish (*Gasterosteus aculeatus*). Threespine stickleback exhibit a large degree of within and between population genetic and phenotypic variation making them extremely useful for genetically mapping complex traits including differences in gastrointestinal inflammation (Colosimo et al. 2004; Cresko et al. 2004; Hohenlohe et al. 2010; Glazer et al. 2015; Greenwood et al. 2015; Beck et al. 2020). Threespine stickleback have also been developed as a model for understanding genetic and environmental contributions to gut microbiome composition through work in wild populations, controlled laboratory studies enabling environmental manipulations, and gnotobiotic studies (Smith et al. 2015; Milligan-McClellan et al. 2016; Small et al. 2017, 2023; Rennison et al. 2019; Steury et al. 2019; Fuess et al. 2021). We took advantage of these features of threespine stickleback fish and describe here the phenotypic presentation of these spontaneous mutants exhibiting markers of an endosome trafficking disorder including oculocutaneous albinism and vision impairment and provide evidence of alterations to the gut microbiome based on 16S amplicon sequencing mimicking microbial community shifts common in IBD.

## MATERIALS AND METHODS

### Fish Husbandry

We generated multiple families of threespine stickleback fish (*Gasterosteus aculeatus*) using an F1-intercross framework originating from different parental populations. This included females from Rabbit Slough, Alaska (N 61.5595, W149.2583) and males from Eel Creek, Oregon (N 43.5771, W124.1932). We performed all crosses via *in vitro* fertilization (Cresko et al. 2004) with testes stored in Ringer’s solution with 10mg/mL Gentamycin and 100x concentration of cell culture antibiotic/mycotic from Gibco-BRL which allowed for fertilization of multiple egg clutches from a single dam by a single sire. The largest F2 mapping family consisted of 226 fish generated from three egg clutches of a single dam. We raised these fish in the University of Oregon stickleback aquatic facility where fish were fed hatched brine shrimp nauplii and fry food (Ziegler AP100 larval food) until 60 days post fertilization (dpf). We generated additional adult and juvenile fish, used for imaging, histology, and microbiome analyses in separate F1 intercrosses from the same pool of F1 siblings as the F2 mapping cross. We also selected mutant F2s randomly and crossed them to generate F3 and subsequently F4 generations follow the same *in vitro* fertilization methods (Cresko et al. 2004). We euthanized all fish in MS222 following IACUC approved methods.

### Dissections and Sample Preparation

For genetic mapping, we obtained caudal fin clips - post-euthanasia - from 190 fish which were scored based on presence or absence of pigment. We also obtained caudal fin clips from the F1 parents. For microbiome analyses, we used sex-balanced wildtype adults from both parental populations (n = 6 respectively) and mutant adults from F2 families (n = 38). We dissected the stomach, cutting at the connection between the stomach and intestine. Gut samples were flash frozen in liquid nitrogen and stored at −80°C. We determined sex of individuals for the microbiome study morphologically during the dissection process.

### DNA isolation and sex determination

For genetic mapping, we isolated genomic DNA from the 190 caudal fin clips using the DNeasy Blood and Tissue kit (Qiagen, Valenica, CA, USA) and quantified DNA using the Qubit Fluorometer Broad Range kit (Thermo Fisher Scientific, Waltham, MA USA). We determined the sex of the juvenile fish (60 dpf) using sex specific PCR amplification via the GA1 primer pair (Griffiths et al. 2000) to identify the presence of a male-specific amplicon. We then standardized genomic DNA to 10ng/μL for Restriction-site Associated DNA (RAD) sequencing library preparation following previously published methods (Baird et al. 2008; Hohenlohe et al. 2010; Etter et al. 2012). For microbiome analyses, we homogenized guts following a two-step homogenization pipeline and isolated DNA following modified protocols (Small et al. 2019) using Qiagen DNeasy spin columns (Qiagen, Valenica, CA, USA). We quantified DNA using the Qubit Fluorometer Broad Range kit (Thermo Fisher Scientific, Waltham, MA USA) and standardized to 20ng/μL for 16S rRNA (V4 region) sequencing library preparation.

### Library Preparation and Sequencing

To generate RAD-seq libraries for genetic mapping, we digested genomic DNA using the restriction endonuclease SbfI-HF (New England Biolabs, Ipswitch, MA, USA). We uniquely barcoded samples and multiplexed them into a single lane of sequencing on an Illumina HiSeq 4000 to obtain single-end 150 nucleotide (nt) reads. To generate 16S-sequencing libraries, we submitted genomic DNA isolated from guts to the University of Oregon Genomics and Cell Characterization Core Facility (GC3F) for V4 region amplicon library preparation. This was followed by individual barcoding, multiplexing, and sequencing on an Illumina HiSeq 4000 to obtain paired-end 150 nt (nucleotide) reads.

### Genetic Mapping

We demultiplexed raw reads using the process_radtags program in the Stacks suite v1.48 (Catchen et al. 2011, 2013). We aligned reads using GSNAP (Wu and Nacu 2010) to the threespine stickleback reference genome from Ensembl (v80) allowing seven maximum mismatches and a terminal threshold of ten to avoid soft-clipping and spurious alignment of short reads. We converted Sam to Bam using VCFtools (Danecek et al. 2011) and sorted Bam files using samtools v 1.19 (Li et al. 2009). To build loci and call genotypes for all individuals we used the gstacks program from the Stacks v2.68 suite (Catchen et al. 2011, 2013; Rochette et al. 2019) with default settings, except that we set the *min-mapq* flag to 10 and the *max-clipped* flag to 0.2. Using the populations program within Stacks we generated onemap and rqtl files, which we then converted to so-called “raw” and “genetic map” files required for mapping using R/qtl2 (Broman et al. 2019), with a custom shell script. Within the populations program we used default options, except that we set the *min-populations* flag to 2, the *min-samples-per-pop* flag to 0.5, the *max-obs-het* flag to 0.5, the *map-type* flag to F2, and we enabled the *filter-haplotype-wise* flag. To perform genetic mapping, we assigned phenotypes using a binary score 0 = wildtype and 1 = mutant and used the R/qtl2 *calc_genoprob* function, followed by the *scan1* function with Haley-Knott regression and a binary trait model. We determined significance of associations by conducting permutation tests of 1,000 permutations using the *scan1perm* function.

### Phenotyping, Imaging, and histology

To generate a developmental time series, we used both mutants from an F2 family and wildtype Rabbit Slough fish. We randomly selected five fish from the families every 3 days and fixed them in 4% paraformaldehyde (PFA) at room temperature overnight then moved to 4°C for long-term storage. We then randomly selected a representative from each timepoint for imaging on the Olympus SZX17 Stereo microscope. To capture adult phenotypes impacting eye pigmentation, we collected heads of wildtype Rabbit Slough adults (n = 6) and F2 mutants (n = 6). We cut the head posterior to the gill rakers, preserved it in 4% PFA, and submitted to the University of Oregon histology core. Experts at the histology core sectioned the eyes and stained them using a Hematoxylin and Eosin (H&E) stain. We imaged these sections using the Olympus BX51 compound microscope.

### Microbiome analysis

We joined and filtered raw reads in QIIME2 version 2019.7 (Bolyen et al. 2019) using default parameters for the “vsearch join-pairs plugin” (Rognes et al. 2016) and 175 bp as a minimum read length to exclude stickleback 12 mitochondrial amplicons. We filtered based on quality-scores using the “quality-filter” plugin and default parameters (Bokulich et al. 2013) and performed denoising and trimming with Deblur (deblur denoise-16S plugin) (Amir et al. 2017) which also generated the ASV table. We assigned taxonomy to feature IDs using a Greengenes (13_8 99% OTUs) classifier (DeSantis et al. 2006; McDonald et al. 2012) trained with 515/806 primers with the “feature-classifier classify-sklearn” plugin (Pedregosa, F. et al. 2011; Bokulich et al. 2018) and used the program “R” (version: 4.3.2, R Core Team 2023) for all subsequent data analyses.

We removed mitochondrial and chloroplast feature IDs from the taxonomy table and ASV table. These feature IDs accounted for 1186 ASVs. We also removed sample 69.16, as there were problems that arose during removal of the gut including dissection notes of a twisted gut that heavily impacted morphology. We measured alpha diversity using two metrics: observed ASV richness and Pielou’s Evenness (Pielou 1966) using the function ‘lme’ from the R package ‘nlme’ (Pinheiro et al. 1999; Pinhero, J.C. and Bates, D.M. 2000) with ‘Tank’ as a random effect to test if alpha diversity varied between mutant and wildtype stickleback. We measured beta-diversity using Bray-Curtis Dissimilarity (Bray and Curtis 1957) and used the ‘adonis2’ function from the ‘vegan’ package (Oksanen et al. 2001) to run a PERMANOVA (99,999 iterations) with the distance matrix as a response variable and ‘pigment’ and ‘tank’ as independent variables. To determine differentially abundant bacterial phyla between mutant and wildtype stickleback, we used a generalized linear mixed model (GLMM), which has been used previously to address differentially abundant taxa NBZIMM (Zhang and Yi 2020). We clustered ASVs to the phylum level and tested the differential abundance of each phylum independently. Using the R package ‘glmmTMB’ (Brooks et al. 2017) we ran a GLMM with negative binomial distribution as the family, the phylum in question as the response variable and pigment as the independent variable. We supplied the natural log of the total reads per sample as an offset term to account for differing sampling depths. Tank and Sample ID were used as random effects in the models. We tested each model for dispersion and zero-inflation using the package ‘DHARMa’ (Hartig 2016) and use the model.sel function from the package ‘MuMIn’ (Bartoń 2010) to identify best fit models based on lowest AIC for each phylum. We then adjusted the p-values for false discovery rate using the False Discovery Rate (Benjamini and Hochberg 1995).

## RESULTS AND DISCUSSION

### Identification and genetic mapping of a pigment mutant supports a single gene mutation

In a family of 226 threespine stickleback fish, 57 appeared albino, yielding a 25.2% mutant phenotype frequency indicative of a single gene mutation (Fig. S1). To test our single-gene hypothesis, we crossed mutant F2s together which yielded 100% mutant offspring. We repeated this again in the next generation, intercrossing F3s, and again 100% of offspring appeared mutant. Combined, these data support our hypothesis of a single-gene mutation causing a loss of melanin. To investigate the specific genetic mutation underpinning the mutant phenotype we took advantage of the already generated F2 mapping family. Using RAD-seq we genotyped 190 F2s (including 57 mutants) as well as the F1 parents. We then used genetic mapping to identify genomic regions strongly associated with the loss of melanin (Fig. 1). We identified a strong genomic association with loss of pigment – not surprising given our single-gene mutation hypothesis. We specifically observed a significant association with chromosome 13. This chromosome contained multiple significant peaks. When exploring the full chromosome, we identified 11 genes with KEGG (Kanehisa 2000) associations to endosome trafficking (Table 1), however we did not find a causative lesion. Despite not finding the specific mutation, the phenotypic analysis is informative for the mode of action.

**Figure 1.**
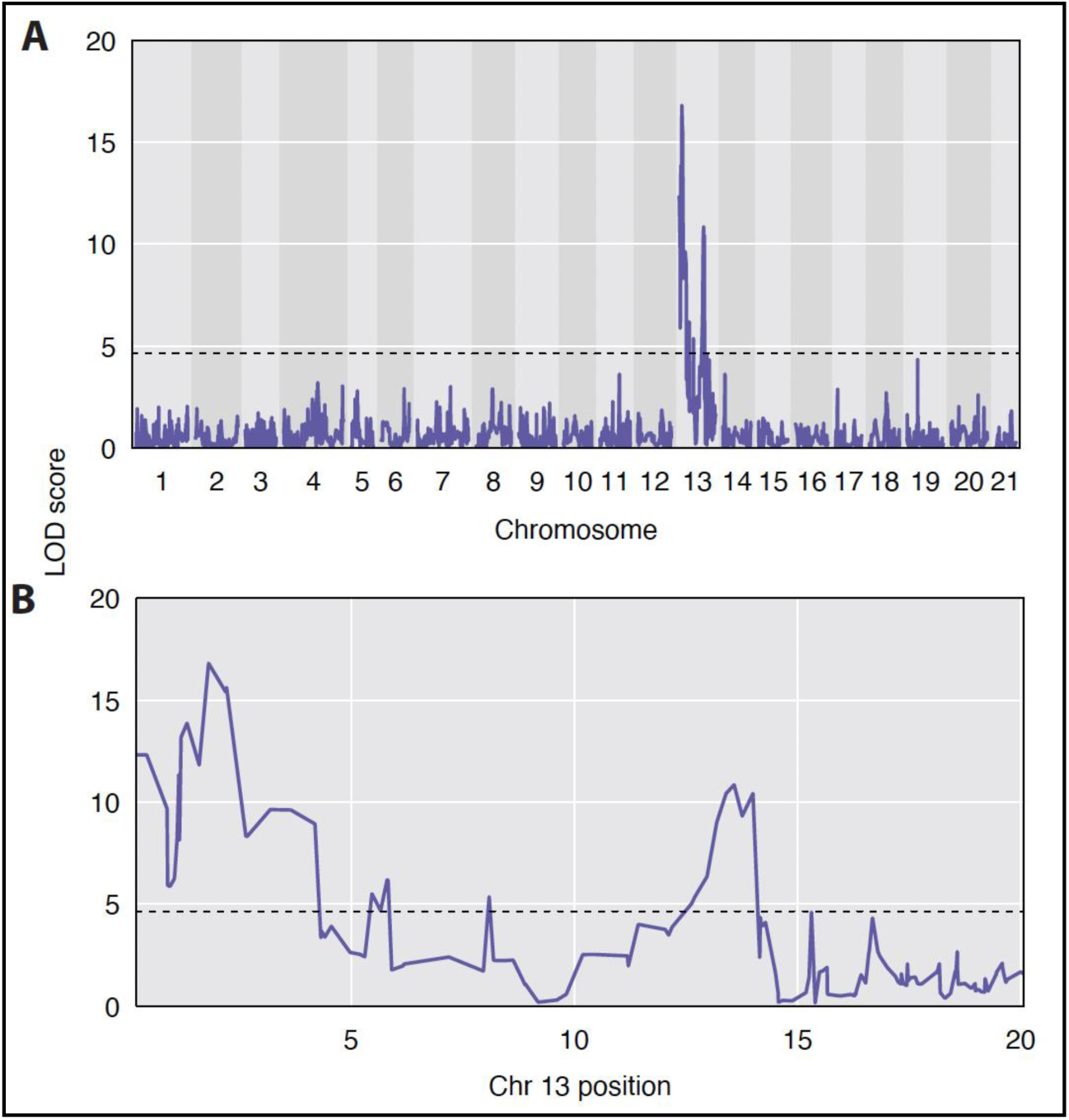
QTL Mapping of pigment loss phenotype. LOD (logarithm of the odds) indicates strength of association. Background shading differences indicate chromosome boundaries. Horizontal dashed line indicates significance threshold determined by a permutation test of 1,000 permutations. (A) Genome wide view of association strength. (B) View of Chromosome 13 only. Position given in Mb.

**Table 1.** KEGG associated endosome trafficking genes on LG13 in threespine stickleback.

| Name | Genomic Position | Endosome-Lysosome Transport Pathway |
| --- | --- | --- |
| GOLPH3 | groupXIII:1,388,883-1,391,798 | Rab Associated Protein (Rab GTPases) |
| VPS37B | groupXIII:2,950,458-2,959,985 | Endosomal Sorting Complexes (ESCRT-I) |
| AP3S1 | groupXIII:4,627,086-4,635,678 | Adaptor Complex Protein (AP-3) |
| NAA35 | groupXIII:7,397,696-7,405,305 | N-acetyl Transferase Complex C (NatC) |
| RILPL1 | groupXIII:8,293,913-8,298,787 | Rab Associated Protein (Rab GTPases) |
| VPS33A | groupXIII:12,875,214-12,880,198 | Tethering Complex (CORVET/Hops) |
| MVB12BB | groupXIII:13,673,988-13,697,933 | Endosomal Sorting Complexes (ESCRT-I) |
| CHMP7 | groupXIII:14,423,967-14,427,196 | Endosomal Sorting Complexes (ESCRT-III) |
| HPS4 | groupXIII:14,717,817-14,721,920 | Biogenesis LRO Complex (BLOC-3) |
| AP3M2 | groupXIII:15,975,063-15,980,229 | Adaptor Complex Protein (AP-3) |
| STAMBP | groupXIII:18,916,597-18,920,696 | STAM binding protein (Other) |

### Mutants exhibit a loss of melanophores in healthy melanocytes suggesting impaired endosome trafficking

In these spontaneous mutants, we identified a complete loss of melanin leading to oculocutaneous albinism. There was some variation in the severity of the pigment-loss phenotype, but all mutant fish exhibited severe loss of melanin-based pigmentation (Fig. S1). We did not observe any association of phenotype with sex suggesting the mutation is not sex-linked. This is consistent with our mapping results, as we did not identify any associations on chromosome 19 – the location of the stickleback sex locus (Peichel et al. 2004).

To investigate the source of the melanin loss, we generated a developmental time series of mutant and wildtype fish (Fig. 2). We began this series after hatching, at 10 days post fertilization (dpf) and imaged a random representative fish from mutant or wildtype families every 3 days until 31 dpf (Fig. 2A1-Fig. 2H4). From 10dpf to 19dpf we observed a small amount of melanin in the bodies of mutants, noticeably reducing at each 3-day window. After 19dpf, no more melanin was visible in the bodies of albino fish. In the eye, however, we noticed a persistence of dark pigmentation which was not lost until adulthood (Fig. S1).

**Figure 2.**
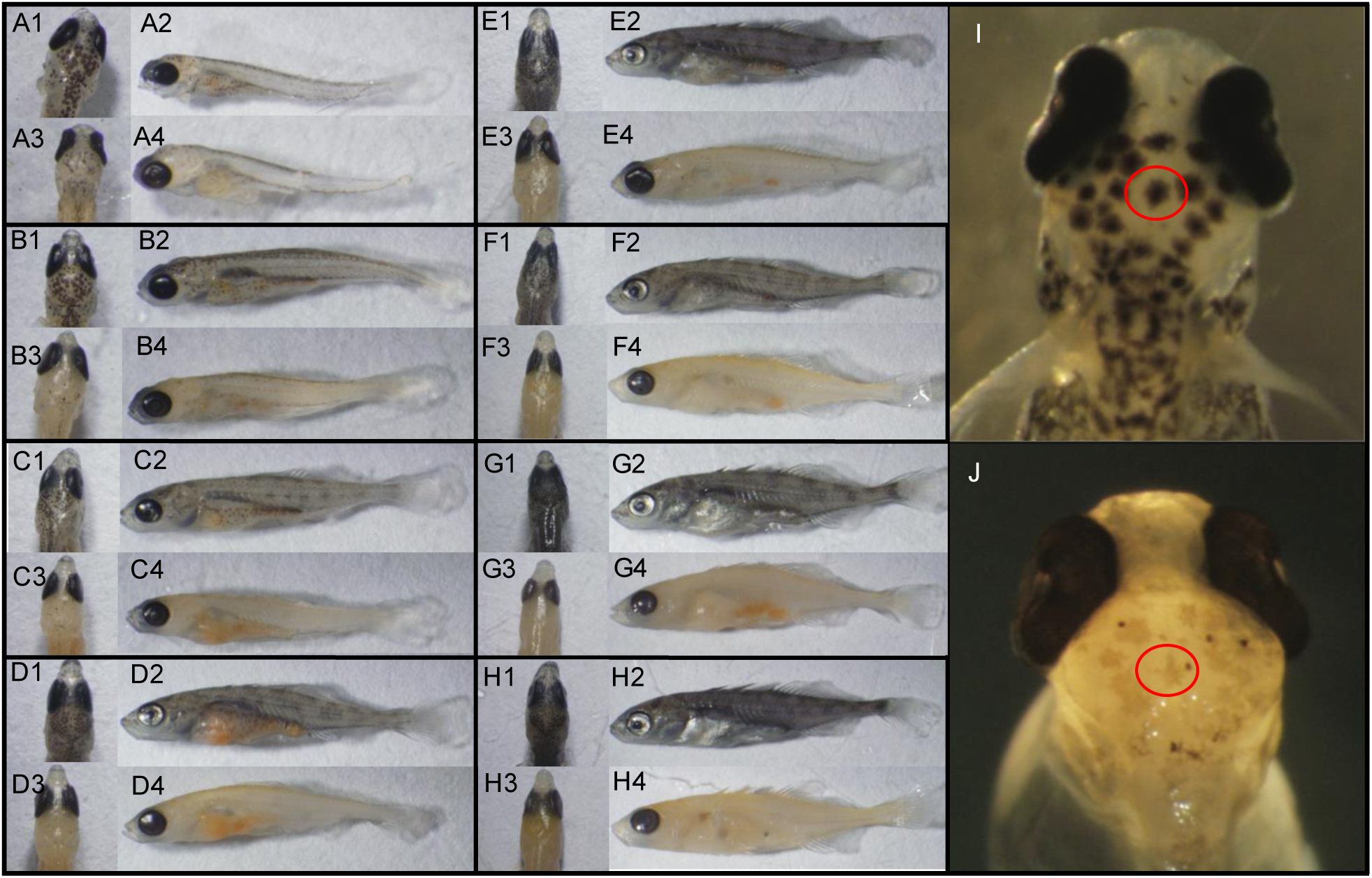
Developmental time series and imaging of melanophores and intact melanocytes in mutant vs wildtype fish. For each panel group A-H a single timepoint is shown comparing wildtype fish (top row) to mutant fish (bottom row). Dorsal views of the head are shown on the left and full body lateral views on the right. (A) 10 days post fertilization ( dpf), (B) 13 dpf, (C) 16dpf, (D) 19 dpf, (E), 22dpf, (F) 25 dpf, (G) 28dpf, (H) 31 dpf. Panels I and J show normal melanocyte morphology in wildtype (I) with lysed melanosomes in mutants (J). Red circle indicates a single representative melanocyte in each fish.

To identify the mechanism of loss of pigment – i.e., deficiency in melanophores vs. melanocytes – we imaged melanocytes prior to complete loss of melanin. In mutant fish, melanocytes are clearly visible and appear wildtype in morphology, however intact organelles containing the pigment (melanophores) are severely lacking. In mutants, a grey tint is visible throughout the mutant fish melanocytes with only a few intact melanophores containing melanin (Fig. 2I; Fig. 2J). The loss and lysis of melanophores over time suggests a failure of melanophores to mature, with organelles lysing and leaking melanin leading to the grey tint and is suggestive of a mutation in endosome trafficking (Raposo and Marks 2007). We further observed that chromatophores – including iridophores – were not negatively impacted consistent with findings in tilapia with perturbed endosome trafficking (Wang et al. 2022) (Fig. S1).

As juveniles, mutant fish did not fare well when co-housed with wildtype siblings. We hypothesized that they might not be competing well for food and observed that they were often found on the bottom of the tank. We investigated the structures of the eyes to test if pigment loss in the eye was impacting vision. To this end, we compared H&E-stained sections of adult eyes from mutant and wildtype fish. Most striking and repeatable was the loss of the retinal pigment epithelium (RPE) a protective melanin-pigmented layer essential for survival of photoreceptors and proper retinal development (Iwai-Takekoshi et al. 2016) (Fig. 3). Not surprisingly, we noticed that the loss of the RPE led to destruction of the photoreceptor layer containing rods and cones, and severe reduction of the plexiform layers commonly associated with vision loss (Moster et al. 2016) (Fig 3D–Fig 3F). In some sections, invagination of additional eye layers was also present (Fig 3F) indicating a likely disruption to vision. Combined, these results suggest that in addition to developing oculocutaneous albinism our fish also exhibit vision lost – both common symptoms of human suffering from endosome trafficking disorders.

**Figure 3.**
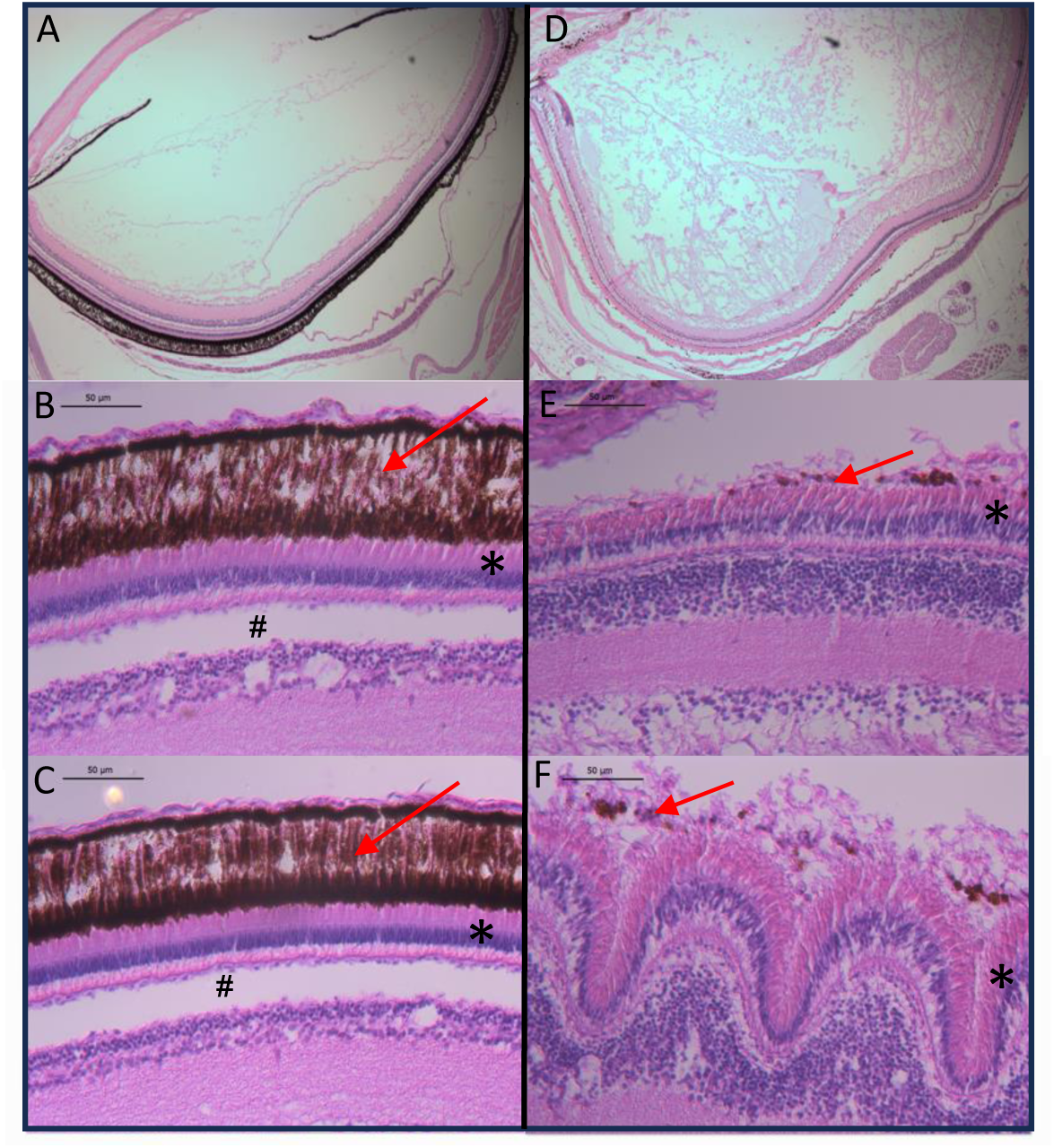
Cross section imaging of eye layers in wildtype and mutant fish. Wildtype H &E stained sections are shown on the left and mutant sections on the right. (A & D) show a full lens cross section. (B &C) show limited variation in retinal pigment epithelial (RPE) layers in wildtype fish (E & F) show substantial structural variation in mutant fish. Red arrows indicate pigment layer, * - indicates the rod and cone layer, # - indicates the plexiform layer (not present in mutant fish).

### Mutations in endosome trafficking impact the intestinal microbiome mimicking IBD

In addition to oculocutaneous albinism and vision loss, humans suffering from endosome trafficking disorders also exhibit intestinal symptomologies consistent with Inflammatory Bowel Diseases (IBD) (Schinella et al. 1980; Erzin et al. 2006; Grucela et al. 2006; Hazzan et al. 2006; Leusse et al. 2006; Wei 2006; Kouklakis et al. 2007). Some endosome trafficking animal mutants are also used to study IBD due to similar symptom presentation (Itoh et al. 2016). While the shared presentation of intestinal inflammation between endosome trafficking disorders and IBDs is well established, the possible shared impacts on the gut microbiome was unexplored. We investigated if changes to the microbiome in our endosome trafficking mutants were consistent with those of IBD models. We obtained 16S-sequencing data from adult mutant and wildtype guts. We included wildtype individuals representing both originating populations of the mutant F2s to account for population-level influences on the microbiome (Small et al. 2023). Our hypothesis was that inflammation would lead to a reduction in alpha diversity and cause specific shifts in abundance of the major phyla of a healthy gut – Firmicutes, Bacteroidetes, Proteobacteria, and Actinobacteria – in directions mimicking inflammation (Matsuoka and Kanai 2015; Wright et al. 2015). More specifically, we expected to observe a decrease in Firmicutes (Chen et al. 2014; Wright et al. 2015), an increase in Proteobacteria and Actinobacteria, and a change in Bacteroidetes (as shifts in both directions have been reported in Crohn’s disease and Ulcerative Colitis patients) (Chen et al. 2014; Matsuoka and Kanai 2015; Wright et al. 2015).

First, we investigated total patterns of alpha and beta diversity between the microbiomes of wildtype individuals and mutant individuals. We found no differences in species richness (estimate = −21.717, t = −0.22, p = 0.840) or evenness (estimate = −0.005009, t = −0.152, p = 0.88) between wildtype and mutant individuals (Fig. S2). However, pigment phenotype (mutant vs wildtype) significantly explained variation in microbiome composition (Bray-Curtis, *PERMANOVA*, R2 = 0.04697, F = 2.606, p = 0.013), although tank explained more variation (Bray-Curtis, *PERMANOVA*, R2 = 0.16, F = 2.96, p < 0.00001) (Fig. S2).

Next, we tested for differential abundance of phyla between the mutant and wildtype groups and observed patterns consistent with an inflamed gut (Fig 4). Specifically, we identified a significant decrease in abundance of Firmicutes in mutant vs wildtype (β = 1.65, SE = 0.645, z = 2.55, p_FDR_ = 0.0203), a significant increase in Proteobacteria (β = −0.402, SE = 0.0835, z = −4.81, p_FDR_ = 2.88×10^−6^), and suggestive patterns of a shift in Bacteroidetes to more abundant in mutant fish (β = −0.518, SE = 0.312, z = −1.66, p_FDR_ = 0.163), although this result was not statistically significant. We did not observe any changes in Actinobacteria as predicted (β = 0.148, SE = 0.665, z = 0.222, p_FDR_ = 0.935) though there is some variation is the shifts of certain groups of Actinobacteria (Oka and Sartor 2020) and more inclusive metagenomic analyses could help disentangle shifts further. We did however observe other changes in phyla not comprising a typical core gut microbiome including a decrease in TM7 in mutant fish (β = 1.91, SE = 0.815, z = 2.34, p_FDR_ = 0.0360). Previous studies found that patients with Crohn’s disease had a higher diversity of TM7 (Kuehbacher et al. 2008). However, we did not detect differences in richness or evenness of TM7 between the wildtype and mutant individuals (Fig S3). While not statistically significant, Acidobacteria were trending toward more abundant in mutant fish (β = −0.754, SE = 0.362, z = −2.08, p_FDR_ = 0.0679). This is similar to previous findings that Acidobacteria is enriched in patients with Crohn’s disease (Chen et al. 2014). The patterns observed in the gut microbiota of our mutants suggest shifts in phyla are consistent between individuals with IBD and endosome trafficking disorders possibly connecting these two groups of diseases beyond the current connection of chronic inflammation.

**Figure 4.**
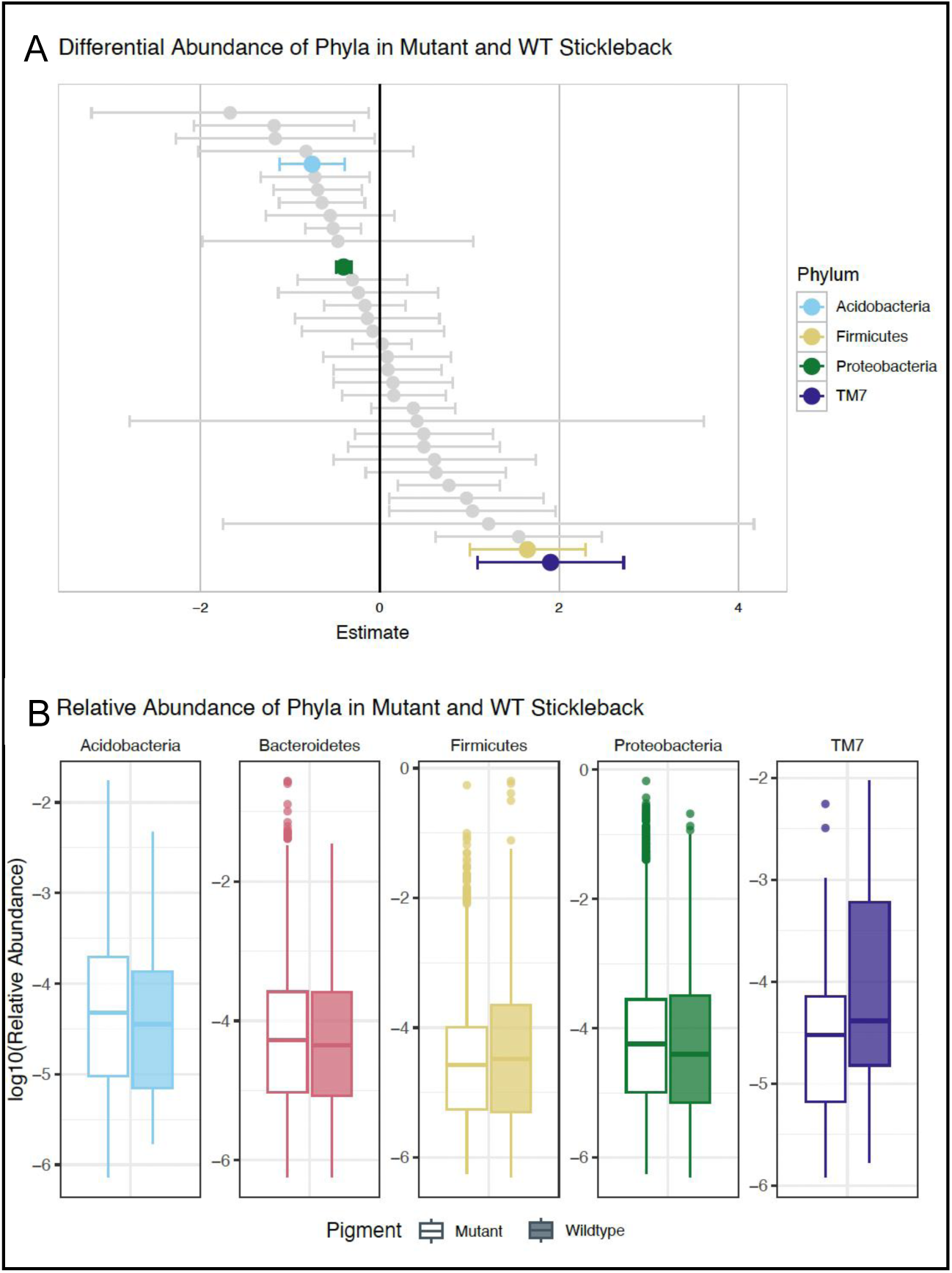
Analysis of microbiome shifts in wildtype vs mutant fish. A) Differential abundance estimates were calculated using a general linear mixed model (GLMM). Positive estimates represent phyla that are more abundant in wildtype stickleback than mutant stickleback. Negative estimates represent the opposite. The width of the error bars is two times the standard error. Marginally significant phyla (p_FDR_ <= 0.1) are highlighted. B) Boxplots of log_10_ Relative Abundance of marginally significant phyla (and Bacteroidetes) in mutant and wildtype stickleback.

### Applications of endosome trafficking mutants beyond IBD

Similar shifts in our endosome trafficking mutant gut microbiota to IBD highlight the possible utility of HPS and other endosome trafficking mutants to study IBD. More importantly, connecting host-microbiome shifts between these groups of disorders suggests interventions developed to treat IBD may be effective in combating chronic inflammation observed in endosome trafficking disorders. This is particularly important as many interventions designed to treat extreme cases of enterocolitis are invasive surgical procedures (Pittet et al. 2013) which are contraindicated, but possible with additional intervention in patients with endosome trafficking disorders due to blood clotting deficiencies (Van Dorp et al. 1990; Lederer et al. 2005; El-Chemaly et al. 2018).

Previous groups have identified mutations in similar pathways including disease pathways associated with Hermansky-Pudlak Syndromes in humans. These include a sex-linked *hps5* mutation in threespine stickleback (Hart and Miller 2017) and *hps4* in channel catfish (Li et al. 2017) and tilapia (Wang et al. 2022). Induced mutations in *hps* genes have also been used to study impacts of melanosome reduction on other chromatophores in Tilapia and stickleback with these mutations have been maintained for the purposes of creating GFP reporter lines with increased visibility due to the transparent appearance of fish without melanosomes. We caution the use of endosome trafficking mutants as “transparent reporter lines” due to other phenotypes deviating from wildtype including - but not limited to - intestinal inflammation.

Pervasiveness of endosome trafficking mutants persisting in natural populations including tilapia, channel catfish, and threespine stickleback warrants more investigation. In the case of our mutants, individuals were generated from a recent influx of sperm from males caught in the wild – suggesting these mutations exist at some low frequency in the population. It is possible, that there is some heterozygote advantage maintaining these variants. This warrants further exploration into populations harboring these mutations.

## CONLCUSIONS

Our findings support the identification of a threespine stickleback fish endosome trafficking mutant exhibiting oculocutaneous albinism and vision impairment caused by disrupted melanosome maturation. The gut microbiota of our mutants demonstrate that shifts in phyla in endosome trafficking mutants are consistent with individuals with IBD possibly connecting these two groups of diseases beyond the current connection of chronic inflammation. We argue that these findings suggest a possible use of HPS and other endosome trafficking mutants to study IBD and that interventions developed to treat IBD may be effective in combating chronic inflammation in endosome trafficking disorders. Given the strengths of threespine stickleback fish as a model for exploring the role of natural genomic variants underlying complex traits and the extensive toolkit available for exploration of the gut microbiome, we argue these fish could serve as excellent models for understanding the connection between endosome trafficking and IBD. However, we further contend these research avenues may be fruitful in treatment of not just intestinal inflammation, but pulmonary inflammation and should be explored in other established murine models.

## Acknowledgements

We would like to thank Emily Niebergall for help with imaging for the developmental times series, Hannah Tavalire for input on the microbiome analyses, and Susan Bassham for helpful comments experimental design. We would also like to thank the members of the META Center (Microbial Ecology and Theory of Animals) for guidance throughout the study and during manuscript preparation especially Karen Guillemin.

## Author Contributions

The study was designed by E.A.B. and W.A.C. Endosome phenotyping was performed by E.A.B. Microbiome analyses were performed by C.A.S. with input from B.J.B and W.A.C. Genetic mapping was performed by C.M.S and E.A.B. RAD-seq libraries were made by M.C.C. The manuscript was primarily written by E.A.B. and C.A.S with all authors contributing to subsequent versions.

---

**Figure S1.**
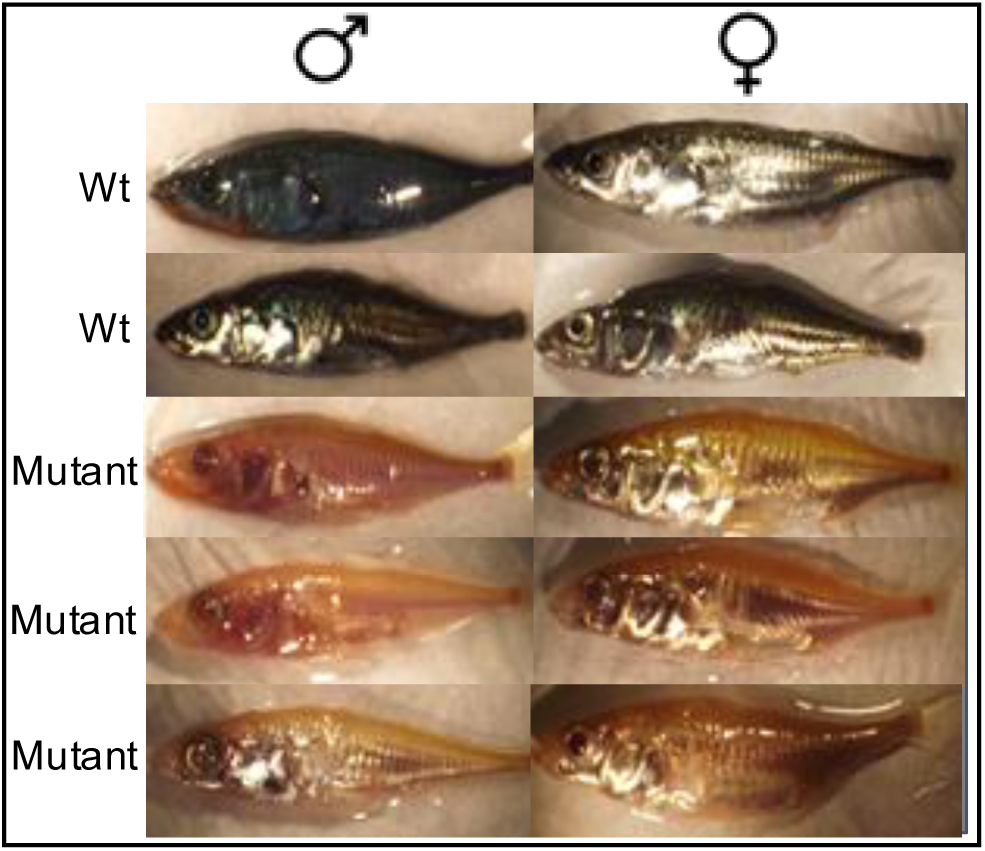
Phenotypic variation by sex of wildtype (Wt) vs Mutant.

**Fig S2:**
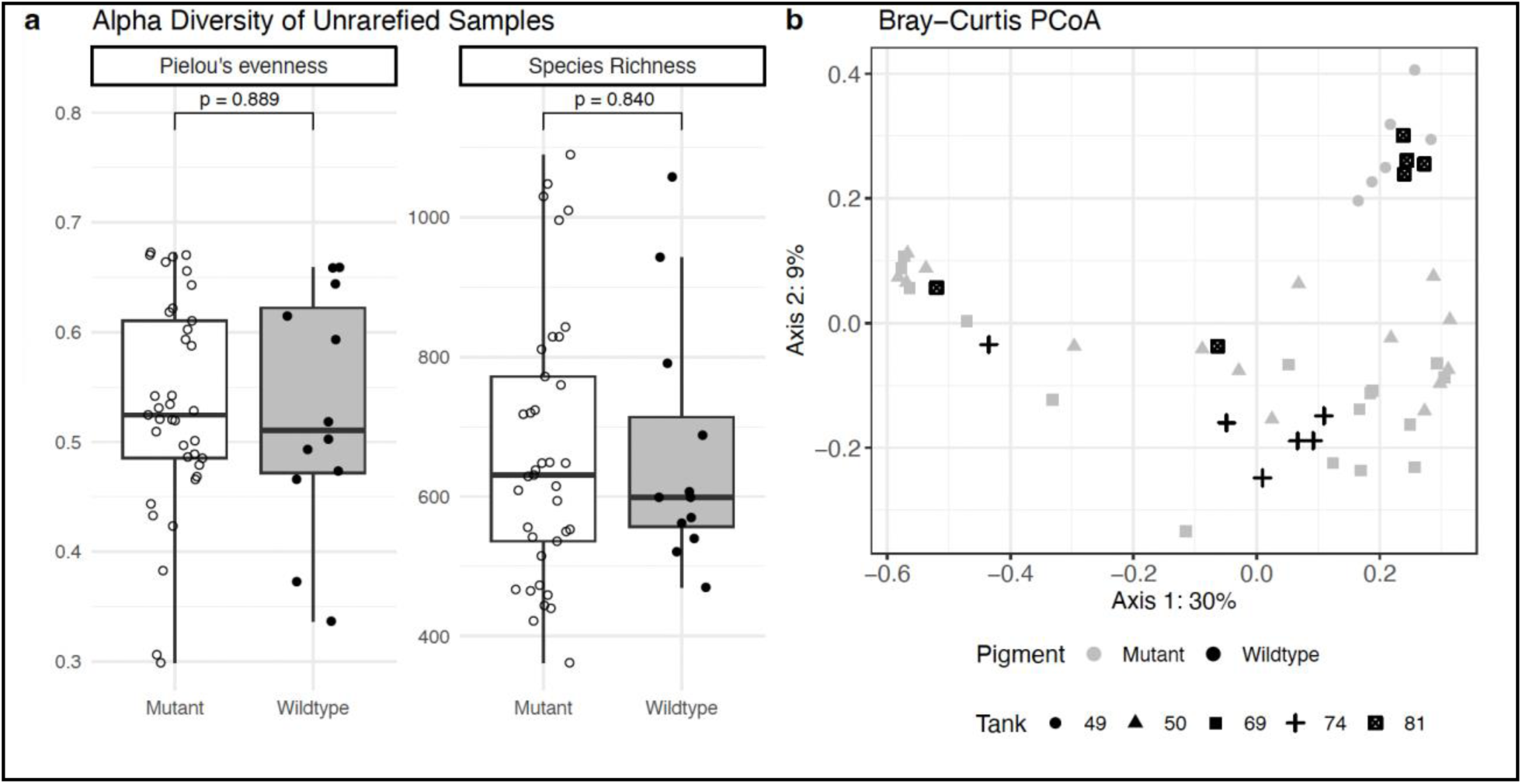
A) Alpha Diversity measured with Pielou’s Evenness and Species (ASV) Richness. Differences in alpha diversity between mutant and wildtype stickleback were tested using a linear effects model. No significant differences were detected between mutant and wildtype stickleback. B) Principal Coordinates Analysis of Bray Curtis Dissimilarity between stickleback reveal minor compositional differences in the microbiomes of mutant and wildtype stickleback.

## References

Amir, A., D. McDonald, J. A. Navas-Molina, E. Kopylova, J. T. Morton, Z. Zech Xu, E. P. Kightley, L. R. Thompson, E. R. Hyde, A. Gonzalez, and R. Knight. 2017. Deblur Rapidly Resolves Single-Nucleotide Community Sequence Patterns. mSystems 2:e00191–16.

Baird, N. A., P. D. Etter, T. S. Atwood, M. C. Currey, A. L. Shiver, Z. A. Lewis, E. U. Selker, W. A. Cresko, and E. A. Johnson. 2008. Rapid SNP Discovery and Genetic Mapping Using Sequenced RAD Markers. PLoS ONE 3:e3376.

Bartoń, K. 2010.MuMIn: Multi-Model Inference.

Beck, E. A., M. C. Currey, C. M. Small, and W. A. Cresko. 2020. QTL Mapping of Intestinal Neutrophil Variation in Threespine Stickleback Reveals Possible Gene Targets Connecting Intestinal Inflammation and Systemic Health. G3 Genes|Genomes|Genetics 10:613–622.

Benjamini, Y., and Y. Hochberg. 1995. Controlling the False Discovery Rate: A Practical and Powerful Approach to Multiple Testing. Journal of the Royal Statistical Society Series B: Statistical Methodology 57:289–300.

Bokulich, N. A., B. D. Kaehler, J. R. Rideout, M. Dillon, E. Bolyen, R. Knight, G. A. Huttley, and J. Gregory Caporaso. 2018. Optimizing taxonomic classification of marker-gene amplicon sequences with QIIME 2’s q2-feature-classifier plugin. Microbiome 6:90.

Bokulich, N. A., S. Subramanian, J. J. Faith, D. Gevers, J. I. Gordon, R. Knight, D. A. Mills, and J. G. Caporaso. 2013. Quality-filtering vastly improves diversity estimates from Illumina amplicon sequencing. Nat Methods 10:57–59.

Bolyen, E., J. R. Rideout, M. R. Dillon, N. A. Bokulich, C. C. Abnet, G. A. Al-Ghalith, H. Alexander, E. J. Alm, M. Arumugam, F. Asnicar, Y. Bai, J. E. Bisanz, K. Bittinger, A. Brejnrod, C. J. Brislawn, C. T. Brown, B. J. Callahan, A. M. Caraballo-Rodríguez, J. Chase, E. K. Cope, R. Da Silva, C. Diener, P. C. Dorrestein, G. M. Douglas, D. M. Durall, C. Duvallet, C. F. Edwardson, M. Ernst, M. Estaki, J. Fouquier, J. M. Gauglitz, S. M. Gibbons, D. L. Gibson, A. Gonzalez, K. Gorlick, J. Guo, B. Hillmann, S. Holmes, H. Holste, C. Huttenhower, G. A. Huttley, S. Janssen, A. K. Jarmusch, L. Jiang, B. D. Kaehler, K. B. Kang, C. R. Keefe, P. Keim, S. T. Kelley, D. Knights, I. Koester, T. Kosciolek, J. Kreps, M. G. I. Langille, J. Lee, R. Ley, Y.-X. Liu, E. Loftfield, C. Lozupone, M. Maher, C. Marotz, B. D. Martin, D. McDonald, L. J. McIver, A. V. Melnik, J. L. Metcalf, S. C. Morgan, J. T. Morton, A. T. Naimey, J. A. Navas-Molina, L. F. Nothias, S. B. Orchanian, T. Pearson, S. L. Peoples, D. Petras, M. L. Preuss, E. Pruesse, L. B. Rasmussen, A. Rivers, M. S. Robeson, P. Rosenthal, N. Segata, M. Shaffer, A. Shiffer, R. Sinha, S. J. Song, J. R. Spear, A. D. Swafford, L. R. Thompson, P. J. Torres, P. Trinh, A. Tripathi, P. J. Turnbaugh, S. Ul-Hasan, J. J. J. Van Der Hooft, F. Vargas, Y. Vázquez-Baeza, E. Vogtmann, M. Von Hippel, W. Walters, Y. Wan, M. Wang, J. Warren, K. C. Weber, C. H. D. Williamson, A. D. Willis, Z. Z. Xu, J. R. Zaneveld, Y. Zhang, Q. Zhu, R. Knight, and J. G. Caporaso. 2019. Reproducible, interactive, scalable and extensible microbiome data science using QIIME 2. Nat Biotechnol 37:852–857.

Bowman, S. L., J. Bi-Karchin, L. Le, and M. S. Marks. 2019. The road to lysosome-related organelles: Insights from Hermansky-Pudlak syndrome and other rare diseases. Traffic 20:404–435.

Bray, J. R., and J. T. Curtis. 1957. An Ordination of the Upland Forest Communities of Southern Wisconsin. Ecological Monographs 27:325–349.

Broman, K. W., D. M. Gatti, P. Simecek, N. A. Furlotte, P. Prins, Ś. Sen, B. S. Yandell, and G. A. Churchill. 2019. R/qtl2: Software for Mapping Quantitative Trait Loci with High-Dimensional Data and Multiparent Populations. Genetics 211:495–502.

Brooks, M., E., K. Kristensen, K. Benthem J., van, A. Magnusson, C. Berg W., A. Nielsen, H. Skaug J., M. Mächler, and B. Bolker M. 2017. glmmTMB Balances Speed and Flexibility Among Packages for Zero-inflated Generalized Linear Mixed Modeling. The R Journal 9:378.

Catchen, J., P. A. Hohenlohe, S. Bassham, A. Amores, and W. A. Cresko. 2013. Stacks: an analysis tool set for population genomics. Mol Ecol 22:3124–3140.

Catchen, J. M., A. Amores, P. Hohenlohe, W. Cresko, and J. H. Postlethwait. 2011. *Stacks* : Building and Genotyping Loci *De Novo* From Short-Read Sequences. G3 Genes|Genomes|Genetics 1:171–182.

Chen, L., W. Wang, R. Zhou, S. C. Ng, J. Li, M. Huang, F. Zhou, X. Wang, B. Shen, M. A. Kamm, K. Wu, and B. Xia. 2014. Characteristics of Fecal and Mucosa-Associated Microbiota in Chinese Patients With Inflammatory Bowel Disease. Medicine 93:e51.

Chung, H., S. J. Pamp, J. A. Hill, N. K. Surana, S. M. Edelman, E. B. Troy, N. C. Reading, E. J. Villablanca, S. Wang, J. R. Mora, Y. Umesaki, D. Mathis, C. Benoist, D. A. Relman, and D. L. Kasper. 2012. Gut Immune Maturation Depends on Colonization with a Host-Specific Microbiota. Cell 149:1578–1593.

Colosimo, P. F., C. L. Peichel, K. Nereng, B. K. Blackman, M. D. Shapiro, D. Schluter, and D. M. Kingsley. 2004. The Genetic Architecture of Parallel Armor Plate Reduction in Threespine Sticklebacks. PLoS Biol 2:e109.

Cresko, W. A., A. Amores, C. Wilson, J. Murphy, M. Currey, P. Phillips, M. A. Bell, C. B. Kimmel, and J. H. Postlethwait. 2004. Parallel genetic basis for repeated evolution of armor loss in Alaskan threespine stickleback populations. Proc. Natl. Acad. Sci. U.S.A. 101:6050–6055.

Danecek, P., A. Auton, G. Abecasis, C. A. Albers, E. Banks, M. A. DePristo, R. E. Handsaker, G. Lunter, G. T. Marth, S. T. Sherry, G. McVean, R. Durbin, and 1000 Genomes Project Analysis Group. 2011. The variant call format and VCFtools. Bioinformatics 27:2156–2158.

Dell’Angelica, E. C., V. Shotelersuk, R. C. Aguilar, W. A. Gahl, and J. S. Bonifacino. 1999. Altered Trafficking of Lysosomal Proteins in Hermansky-Pudlak Syndrome Due to Mutations in the β3A Subunit of the AP-3 Adaptor. Molecular Cell 3:11–21.

DeSantis, T. Z., P. Hugenholtz, N. Larsen, M. Rojas, E. L. Brodie, K. Keller, T. Huber, D. Dalevi, P. Hu, and G. L. Andersen. 2006. Greengenes, a Chimera-Checked 16S rRNA Gene Database and Workbench Compatible with ARB. Appl Environ Microbiol 72:5069–5072.

El-Chemaly, S., K. J. O’Brien, S. D. Nathan, G. L. Weinhouse, H. J. Goldberg, J. M. Connors, Y. Cui, T. L. Astor, P. C. Camp, I. O. Rosas, M. Lemma, V. Speransky, M. A. Merideth, W. A. Gahl, and B. R. Gochuico. 2018. Clinical management and outcomes of patients with Hermansky-Pudlak syndrome pulmonary fibrosis evaluated for lung transplantation. PLoS ONE 13:e0194193.

Erzin, Y., S. Cosgun, A. Dobrucali, M. Tasyurekli, S. Erdamar, and M. Tuncer. 2006. Complicated granulomatous colitis in a patient with Hermansky-Pudlak syndrome, successfully treated with infliximab. Acta Gastroenterol Belg 69:213–216.

Etter, P. D., S. Bassham, P. A. Hohenlohe, E. A. Johnson, and W. A. Cresko. 2012. SNP Discovery and Genotyping for Evolutionary Genetics Using RAD Sequencing. Pp. 157–178 *in* V. Orgogozo and M. V. Rockman, eds. Molecular Methods for Evolutionary Genetics. Humana Press, Totowa, NJ.

Fuess, L. E., S. den Haan, F. Ling, J. N. Weber, N. C. Steinel, and D. I. Bolnick. 2021. Immune Gene Expression Covaries with Gut Microbiome Composition in Stickleback. mBio 12:e00145–21.

Glazer, A. M., E. E. Killingbeck, T. Mitros, D. S. Rokhsar, and C. T. Miller. 2015. Genome Assembly Improvement and Mapping Convergently Evolved Skeletal Traits in Sticklebacks with Genotyping-by-Sequencing. G3 Genes|Genomes|Genetics 5:1463–1472.

Greenwood, A. K., R. Ardekani, S. R. McCann, M. E. Dubin, A. Sullivan, S. Bensussen, S. Tavaré, and C. L. Peichel. 2015. Genetic mapping of natural variation in schooling tendency in the threespine stickleback. G3 (Bethesda) 5:761–769.

Griffiths, R., K. L. Orr, A. Adam, and I. Barber. 2000. DNA sex identification in the three-spined stickleback. Journal of Fish Biology 57:1331–1334.

Grucela, A. L., P. Patel, E. Goldstein, R. Palmon, D. B. Sachar, and R. M. Steinhagen. 2006. Granulomatous Enterocolitis Associated with Hermansky-Pudlak Syndrome. Am J Gastroenterol 101:2090–2095.

Hart, J. C., and C. T. Miller. 2017. Sequence-Based Mapping and Genome Editing Reveal Mutations in Stickleback *Hps5* Cause Oculocutaneous Albinism and the *casper* Phenotype. G3 Genes|Genomes|Genetics 7:3123–3131.

Hartig, F. 2016.DHARMa: Residual Diagnostics for Hierarchical (Multi-Level / Mixed) Regression Models.

Hazzan, D., S. Seward, H. Stock, S. Zisman, K. Gabriel, N. Harpaz, and J. J. Bauer. 2006. Crohn’s-like colitis, enterocolitis and perianal disease in Hermansky–Pudlak syndrome. Colorectal Disease 8:539–543.

Hohenlohe, P. A., S. Bassham, P. D. Etter, N. Stiffler, E. A. Johnson, and W. A. Cresko. 2010. Population Genomics of Parallel Adaptation in Threespine Stickleback using Sequenced RAD Tags. PLoS Genet 6:e1000862.

Huizing, M., A. Helip-Wooley, W. Westbroek, M. Gunay-Aygun, and W. A. Gahl. 2008. Disorders of Lysosome-Related Organelle Biogenesis: Clinical and Molecular Genetics. Annu. Rev. Genom. Hum. Genet. 9:359–386.

Hurford, M. T., and C. Sebastiano. 2008. Hermansky-pudlak syndrome: report of a case and review of the literature. Int J Clin Exp Pathol 1:550–554.

Introne, W., M. Huizing, M. Malicdan, K. O’Brien, and W. Gahl. 2023. Hermansky-Pudlak Syndrome. P. in Gene Reviews. University of Washington, University of Washington.

Itoh, Y., Y. Nagaoka, Y. Katakura, H. Kawahara, and H. Takemori. 2016. Simple chronic colitis model using hypopigmented mice with a *Hermansky–Pudlak syndrome 5* gene mutation. Pigment Cell Melanoma Res 29:578–582.

Iwai-Takekoshi, L., A. Ramos, A. Schaler, S. Weinreb, R. Blazeski, and C. Mason. 2016. Retinal pigment epithelial integrity is compromised in the developing albino mouse retina. J Comp Neurol 524:3696–3716.

Jovic, M., M. Sharma, J. Rahajeng, and S. Caplan. 2009. The early endosome: a busy sorting station for proteins at the crossroads. Histology and Histopathology 99–112.

Kanehisa, M. 2000. KEGG: Kyoto Encyclopedia of Genes and Genomes. Nucleic Acids Research 28:27–30.

Kouklakis, G., E. I. Efremidou, M. S. Papageorgiou, E. Pavlidou, K. J. Manolas, and N. Liratzopoulos. 2007. Complicated Crohn’s-like colitis, associated with Hermansky-Pudlak syndrome, treated with Infliximab: a case report and brief review of the literature. J Med Case Rep 1:176.

Kuehbacher, T., A. Rehman, P. Lepage, S. Hellmig, U. R. Fölsch, S. Schreiber, and S. J. Ott. 2008. Intestinal TM7 bacterial phylogenies in active inflammatory bowel disease. Journal of Medical Microbiology 57:1569–1576.

Lederer, D. J., S. M. Kawut, J. R. Sonett, E. Vakiani, S. L. Seward, J. G. White, J. S. Wilt, C. C. Marboe, W. A. Gahl, and S. M. Arcasoy. 2005. Successful Bilateral Lung Transplantation for Pulmonary Fibrosis Associated With the Hermansky-Pudlak Syndrome. The Journal of Heart and Lung Transplantation 24:1697–1699.

Leusse, A. D., E. Dupuy, M. Huizing, C. Danel, G. Meyer, R. Jian, and P. Marteau. 2006. Ileal Crohn’s disease in a woman with Hermansky-Pudlak syndrome. Gastroentérologie Clinique et Biologique 30:621–624.

Li, H., B. Handsaker, A. Wysoker, T. Fennell, J. Ruan, N. Homer, G. Marth, G. Abecasis, R. Durbin, and 1000 Genome Project Data Processing Subgroup. 2009. The Sequence Alignment/Map format and SAMtools. Bioinformatics 25:2078–2079.

Li, Y., X. Geng, L. Bao, A. Elaswad, K. W. Huggins, R. Dunham, and Z. Liu. 2017. A deletion in the Hermansky–Pudlak syndrome 4 (Hps4) gene appears to be responsible for albinism in channel catfish. Mol Genet Genomics 292:663–670.

Marks, M. S., H. F. Heijnen, and G. Raposo. 2013. Lysosome-related organelles: unusual compartments become mainstream. Current Opinion in Cell Biology 25:495–505.

Matsuoka, K., and T. Kanai. 2015. The gut microbiota and inflammatory bowel disease. Semin Immunopathol 37:47–55.

McDonald, D., M. N. Price, J. Goodrich, E. P. Nawrocki, T. Z. DeSantis, A. Probst, G. L. Andersen, R. Knight, and P. Hugenholtz. 2012. An improved Greengenes taxonomy with explicit ranks for ecological and evolutionary analyses of bacteria and archaea. The ISME Journal 6:610–618.

Milligan-McClellan, K., C. M. Small, E. K. Mittge, M. Agarwal, M. Currey, W. A. Cresko, and K. Guillemin. 2016. Innate immune responses to gut microbiota differ between oceanic and freshwater threespine stickleback populations. Dis Model Mech 9:187– 198.

Moster, S. J., M. L. Moster, M. Scannell Bryan, and R. C. Sergott. 2016. Retinal Ganglion Cell and Inner Plexiform Layer Loss Correlate with Visual Acuity Loss in LHON: A Longitudinal, Segmentation OCT Analysis. Invest. Ophthalmol. Vis. Sci. 57:3872.

Naslavsky, N., and S. Caplan. 2018. The enigmatic endosome – sorting the ins and outs of endocytic trafficking. Journal of Cell Science 131:jcs216499.

O’Brien, K. J., X. Parisi, N. R. Shelman, M. A. Merideth, W. J. Introne, T. Heller, W. A. Gahl, M. C. V. Malicdan, and B. R. Gochuico. 2021. Inflammatory bowel disease in Hermansky–Pudlak syndrome: a retrospective single-centre cohort study. J Intern Med 290:129–140.

Oka, A., and R. B. Sartor. 2020. Microbial-Based and Microbial-Targeted Therapies for Inflammatory Bowel Diseases. Dig Dis Sci 65:757–788.

Oksanen, J., G. L. Simpson, F. G. Blanchet, R. Kindt, P. Legendre, P. R. Minchin, R. B. O’Hara, P. Solymos, M. H. H. Stevens, E. Szoecs, H. Wagner, M. Barbour, M. Bedward, B. Bolker, D. Borcard, T. Borman, G. Carvalho, M. Chirico, M. De Caceres, S. Durand, H. B. A. Evangelista, R. FitzJohn, M. Friendly, B. Furneaux, G. Hannigan, M. O. Hill, L. Lahti, C. Martino, D. McGlinn, M.-H. Ouellette, E. Ribeiro Cunha, T. Smith, A. Stier, C. J. F. Ter Braak, and J. Weedon. 2001. vegan: Community Ecology Package.

Pedregosa, F., Varoquaux, G., Gramfort, A., Michel, V., Thirion, B., Grisel, O., Blondel, M., Prettenhofer, P., Weiss, R., Dubourg, V., Vanderplas, J., Passos, A., Cournapeau, D., Brucher, M., Perrot, M., and Duchesnay, E. 2011. Scikit-learn: Machine learning in Python. Journal of Machine Learning Research 12:2825–2830.

Peichel, C. L., J. A. Ross, C. K. Matson, M. Dickson, J. Grimwood, J. Schmutz, R. M. Myers, S. Mori, D. Schluter, and D. M. Kingsley. 2004. The Master Sex-Determination Locus in Threespine Sticklebacks Is on a Nascent Y Chromosome. Current Biology 14:1416–1424.

Pickard, J. M., M. Y. Zeng, R. Caruso, and G. Núñez. 2017. Gut microbiota: Role in pathogen colonization, immune responses, and inflammatory disease. Immunological Reviews 279:70–89.

Pielou, E. C. 1966. The measurement of diversity in different types of biological collections. Journal of Theoretical Biology 13:131–144.

Pinheiro, J., D. Bates, and R Core Team. 1999.nlme: Linear and Nonlinear Mixed Effects Models.

Pinhero, J.C. and Bates, D.M. 2000. Mixed-Effects Models in S and S-PLUS. Springer-Verlag, New York.

Pittet, V., G. Rogler, P. Michetti, N. Fournier, J.-P. Vader, A. Schoepfer, C. Mottet, B. Burnand, F. Froehlich, and for the Swiss Inflammatory Bowel Disease Cohort Study Group (see Appendix). 2013. Penetrating or Stricturing Diseases are the Major Determinants of Time to First and Repeat Resection Surgery in Crohn’s Disease. Digestion 87:212–221.

Raposo, G., and M. S. Marks. 2007. Melanosomes--dark organelles enlighten endosomal membrane transport. Nat Rev Mol Cell Biol 8:786–797.

Rennison, D. J., S. M. Rudman, and D. Schluter. 2019. Parallel changes in gut microbiome composition and function during colonization, local adaptation and ecological speciation. Proc. R. Soc. B. 286:20191911.

Rochette, N. C., A. G. Rivera-Colón, and J. M. Catchen. 2019. Stacks 2: Analytical methods for paired-end sequencing improve RADseq-based population genomics. Molecular Ecology 28:4737–4754.

Rognes, T., T. Flouri, B. Nichols, C. Quince, and F. Mahé. 2016. VSEARCH: a versatile open source tool for metagenomics. PeerJ 4:e2584.

Schinella, R. A., M. A. Greco, B. L. Cobert, L. W. Denmark, and R. P. Cox. 1980. Hermansky-Pudlak Syndrome with Granulomatous Colitis. Ann Intern Med 92:20–23.

Small, C. M., E. A. Beck, M. C. Currey, H. F. Tavalire, S. Bassham, and W. A. Cresko. 2023. Host genomic variation shapes gut microbiome diversity in threespine stickleback fish. mBio 14:e00219–23.

Small, C. M., M. Currey, E. A. Beck, S. Bassham, and W. A. Cresko. 2019. Highly Reproducible 16S Sequencing Facilitates Measurement of Host Genetic Influences on the Stickleback Gut Microbiome. mSystems 4:e00331–19.

Small, C. M., K. Milligan-Myhre, S. Bassham, K. Guillemin, and W. A. Cresko. 2017. Host Genotype and Microbiota Contribute Asymmetrically to Transcriptional Variation in the Threespine Stickleback Gut. Genome Biology and Evolution 9:504–520.

Smith, C. C., L. K. Snowberg, J. Gregory Caporaso, R. Knight, and D. I. Bolnick. 2015. Dietary input of microbes and host genetic variation shape among-population differences in stickleback gut microbiota. ISME J 9:2515–2526.

Steury, Currey, Cresko, and Bohannan. 2019. Population Genetic Divergence and Environment Influence the Gut Microbiome in Oregon Threespine Stickleback. Genes 10:484.

Van Dorp, D. B., P. W. Wijermans, F. Meire, and G. Vrensen. 1990. The Hermansky-Pudlak syndrome: Variable reaction to 1-desamino-8D-arginine vasopressin for correction of the bleeding time. Ophthalmic Paediatrics and Genetics 11:237–244.

Wang, C., T. D. Kocher, B. Lu, J. Xu, and D. Wang. 2022. Knockout of Hermansky-Pudlak Syndrome 4 (hps4) leads to silver-white tilapia lacking melanosomes. Aquaculture 559:738420.

Wei, M. L. 2006. Hermansky–Pudlak syndrome: a disease of protein trafficking and organelle function. Pigment Cell Research 19:19–42.

Wood, L. G., N. Shivappa, B. S. Berthon, P. G. Gibson, and J. R. Hebert. 2015. Dietary inflammatory index is related to asthma risk, lung function and systemic inflammation in asthma. Clin Exp Allergy 45:177–183.

Wright, E. K., M. A. Kamm, S. M. Teo, M. Inouye, J. Wagner, and C. D. Kirkwood. 2015. Recent advances in characterizing the gastrointestinal microbiome in Crohn’s disease: a systematic review. Inflamm Bowel Dis 21:1219–1228.

Wu, T. D., and S. Nacu. 2010. Fast and SNP-tolerant detection of complex variants and splicing in short reads. Bioinformatics 26:873–881.

Yokoyama, T., and B. R. Gochuico. 2021. Hermansky–Pudlak syndrome pulmonary fibrosis: a rare inherited interstitial lung disease. Eur Respir Rev 30:200193.

Zhang, X., and N. Yi. 2020. NBZIMM: negative binomial and zero-inflated mixed models, with application to microbiome/metagenomics data analysis. BMC Bioinformatics 21:488.

